# Age-dependent characterization of the carotid and cerebral artery morphologies in a transgenic mouse model of sickle cell anemia using ultrasound and microcomputed tomography

**DOI:** 10.1101/522748

**Authors:** Christian P. Rivera, Li Li, Shuangyi Cai, Nui Pei, George E. McAlear, Keval Bollavaram, Oluwasanmi V. Aryio, Victor O. Omojola, Hannah Song, Wenchang Tan, Yunlong Huo, Manu O. Platt

## Abstract

Children with sickle cell anemia have elevated stroke risks as well as other arterial complications, but morphological changes to large arteries are not well defined, and the focus has been on the microcirculation where deoxygenation promotes sickling of red blood cells. The goal of this study was to define morphological changes in carotid and cerebral arteries in the Townes transgenic sickle cell mouse model, and to specifically determine anatomical measurement differences in mice homozygous for β-globin S mutation (SS) compared to heterozygous (AS) littermate controls. We used a combination of live imaging with ultrasound and microcomputed tomography (micro-CT) imaging of corrosion casted vessels to quantify arterial dimensions and changes in mice 4, 12, and 24 weeks of age with or without sickle cell anemia. 12 week SS mice had significantly larger common carotid artery diameters than AS mice, and significantly larger diameters in the extracranial and intracranial portions of the internal carotid artery (ICA), determined by ultrasound and micro-CT, respectively. There were also side specific differences between the left and right vessels. There was significant narrowing along ICA length in 12-and 24-week SS mice, decreasing by as much as 70%, such that there was no difference in size between the anterior and middle cerebral arteries, where the ICA terminates, by genotype. Significant narrowing along the length was also measured in the anterior cerebral arteries of 12-and 24-week SS mice, but not AS. Collectively, these findings indicate that sickle cell anemia causes increased arterial dimensions in 12-and 24-week aged mice. We also provide these measurements for the common carotid, internal carotid, anterior cerebral, and middle cerebral arteries for left and right sides, for AS and SS genotypes as a reference for other investigators using *in silico* modeling of arterial complications caused by aging with sickle cell anemia.

## Introduction

Sickle cell disease (SCD) is one of the more prevalent blood disorders worldwide with more than 300,000 new individuals born each year afflicted by the disease. Sickle cell anemia (SCA), specifically, is caused by a missense mutation that substitutes glutamic acid for valine in hemoglobin. This change in molecular structure causes the hemoglobin in red blood cells (RBCs) to polymerize and aggregate under de-oxygenated conditions, leading to deformation and increased rigidity of the cell that cause blockages in the microvasculature that can damage organs and lead to painful episodes known as sickle crisis. SCA also has effects on the large arteries, most notably in the elevated occurrence of strokes in individuals with SCA.

The chance of ischemic stroke is greatly elevated for young individuals occurring in 11% of people between the ages of 2 and 18 [1]. Then, during ages 20-30, the risk of ischemic stroke falls, but is replaced with an increased risk of hemorrhagic stroke. Upon reaching early to mid-30s, the risk of ischemic stroke begins to dominate again in individuals with SCA [2]. Age is an important factor for sickle cell-mediated strokes, and mechanisms behind variations in types of strokes during specific times of life with SCA are unknown. Some clues can be extracted from investigating the vascular morphology. In individuals who have suffered ischemic strokes, cerebral angiograms have shown the presence of stenosis [3-5]. Additionally, autopsies of individuals who have died from sickle-related strokes, show defined markers of intimal hyperplasia [4,6].

In order to obtain to provide insight into underlying structural changes that affect stroke risk in relation to age, mouse models can be used. The murine cerebral circulation displays similarities to humans; the brain is supplied by a pair of vertebral and carotid arteries that branch to form a network resembling the Circle of Willis (CoW) in humans [7]. The Townes sickle cell transgenic mouse model has been beneficial for investigating pulmonary and hematological pathophysiology, with only a few studies on cerebrovascular complications of SCA [8-10].

The goal of this study is to determine how morphology of carotid and cerebral arteries change as the mice age, and any differential effects caused by sickle cell anemia. We use a combination of ultrasound and microcomputed topography (micro-CT), to determine morphometries of carotid and cerebrovascular arteries of Townes mouse model. Both techniques have distinct advantages and disadvantages, which, when combined, can provide a comprehensive morphometric characterization. Ultrasound in living mice allows for longitudinal studies that can be used to obtain arterial dimensions during systole and diastole. However, murine vasculature is much smaller than humans, with vessels approximately 1/10^th^ the size introducing limitations in resolution and access to arteries in the skull. Alternatively, advances in micro-CT imaging procedures for small animal models through the use of perfused contrast agents have enable researchers to investigate structures within a voxel size of 8 μm [11]. For these reasons, we also used micro-CT on corrosion casts of the carotid and cerebral arteries in mice. Casts were created by a plastination method using a radiopaque polymer that hardens in the vessels. This cast was then scanned by micro-CT and reconstructed *in silico* to acquire morphological information of the smaller carotid and cerebral arteries. Casts of the arteries have their own tradeoffs in that it is an endpoint analysis, disallowing longitudinal studies from the same subject. These modalities were used to obtain detailed morphologies of the primary branches of the common carotid artery (CCA), internal carotid artery (ICA), middle cerebral artery (MCA), and anterior cerebral artery (ACA) of mice homozygous for human β globin S, the sickle cell allele (SS) or mice heterozygous for human sickle cell allele and wildtype β globin A allele (AS). Our results suggest sickle cell disease alters the vascular morphology in mice 12 and 24 weeks old, and these age-specific changes may be relevant for age-related risk of types of strokes.

## Methods

### Townes Sickle Cell Transgenic Mice

Townes sickle cell transgenic mice (Jackson Laboratories, Bar Harbor ME) were used for the study. This humanized knock-in model of sickle cell anemia was generated by knocking in the human hemoglobin genes (α, β, and γ), including the β-globin S mutation, into mice null for murine hemoglobin genes [8]. Mice homozygous for β-globin S mutation (SS) were compared to littermate heterozygous (AS) controls. Three different ages were used for the study: 4-weeks, 12-weeks, and 24-weeks, with each age corresponding to a different risk and type of stroke risk in humans with SCA (Table 1). Each age group consisted of 4-7 mice [4 weeks old (n_AS_ = 6, n_SS_ = 12 weeks old (n_AS_ = 7, n_SS_ = 4), 24 week (n_AS_ = 4, n_SS_ = 4)], and all mice were cared for at Peking University (Beijing, China) using standard mouse caring techniques, and ethical protocols were followed based on the Institutional Animal Care and Use Committee of Peking University (Beijing, China).

**Table 1.** Relation of mouse and human ages and types of stroke most likely to occur in humans with sickle cell anemia

| Mouse Age (Weeks) | Approximate Human Age (Years) | Most Likely Type of Stroke |
| --- | --- | --- |
| 4 | 1.5 | Ischemic |
| 12 | 20 | Hemorrhagic |
| 24 | 30 | Ischemic |

### High Frequency Ultrasonography

Ultrasound measurements were performed in all mice using a VEVO 2100 Doppler system (VisualSonics, Toronto, ON, Canada) with a 22-55 Mhz transducer (MS550D, wavelength ~50μm – 2mm), and heart rate was continuously monitored. Hair was then removed from the neck and pectoralis, before ultrasound gel (Aquasonic, Parker Laboratories) was applied to improve signal acquisition. The right carotid bifurcation was visualized in the sagittal plane with B-mode by applying the ultrasound beam perpendicular to the arteries, which were identified based on their depth from the surface of the skin, the direction of flow, and spatial orientation. A 2D recording was taken of the right carotid bifurcation over 2-3 seconds, and then the process was repeated for the left side. Dicom recordings were analyzed in ImageJ to determine arterial diameter during systole and diastole. Diameter was determined as perpendicular distance from the top to bottom of a vessel in the sagittal plane. Two measurements were taken to obtain an average common carotid artery diameter at 0.5 and 1.5mm upstream of the bifurcation. In the extracranial internal carotid artery, only one measurement could be taken 0.5mm downstream of the bifurcation. Three measurements were acquired at each of the above locations to obtain an average across multiple cardiac cycles.

### Preparation for arterial corrosion casting

Following ultrasound measurements, each mouse recuperated for 6-24 hours before being prepared for casting of arteries [12,13]. Briefly, mice were anesthetized with urethane at a dose of 1g/kg, and laparotomy was performed to gain access to the abdominal aorta, where a 24-gauge catheter was used to make a direct puncture. An incision was created in the vena cava to provide an exit point for blood, whereupon a heparinized saline (25 U/mL) was injected through the catheter at a pressure of approximately 100-120 mmHg, the systolic pressure in mice [14,15]. Pressure was monitored using a sphygmomanometer, and the heparinized solution was perfused through the mouse until the effluent at the vena cava became clear. Following exsanguination, a 10% solution of buffered formaldehyde was perfused through the catheter for 15 minutes at a pressure of 80 mmHg to fix the blood vessels. Afterwards, 5 mL of Microfil HV 122 Yellow (Flow Tech, Carver, MA) was manually perfused. Following the procedure, the upper torso of the mouse, including the head, was removed and stored in 10% buffered formaldehyde for 5 days before undergoing decalcification in 10% HCl for 48 hours to remove bone.

### Micro-Computed Tomography (Micro-CT)

Carotid and cerebral artery casts were examined using a high-resolution micro-CT imaging system (Quantum GX, Perkin Elmer). Heads to be scanned were placed prone in a specimen tube (diameter 30.7 mm) with the long axis of the head parallel to the tube’s long axis. Micro-CT images were collected over a 4-mm thick section extending from the nose to the back of the neck, scanning in the transverse plane. Each scan was performed with a beam intensity of 80 microamperes, a beam energy of 70 kV, and an isotropic voxel resolution of 50 μm.

### Image Processing and Morphological Analysis

Micro-CT images were used to create 3D models of the cerebral and carotid arteries for each specimen. Mimics Materialise 15.0 (Leuven, Belgium), was used to manually segment and reconstruct the left and right vasculature, extending from the common carotid arteries to the middle cerebral and anterior cerebral arteries. A minimum threshold was optimized for each specimen, which produced the largest vessel diameter without creating large irregularities on the surface. After a threshold was chosen, a preliminary mesh of the vascular network was created, containing arterial segments of the common carotid artery (CCA), external carotid artery (ECA), internal carotid artery (ICA), anterior cerebral artery (ACA), and middle cerebral artery (MCA). Smoothing algorithms from Vascular Modeling Toolkit *(VMTK)* were applied to remove artifacts created from the reconstruction and re-meshing [16]. The internal carotid artery is divided into two sections: the extracranial internal carotid artery (eICA) and intracranial internal carotid artery (ICA), separated by the bifurcation of the ophthalmic artery (OA) branch. After reconstruction, Mimics was used to calculate morphometric centerline properties along the length of each arterial segment at a resolution of 0.05 mm. These morphometric values include the maximum inscribed internal diameter, perimeter, and cross-sectional area, along the length of each arterial segment. When analyzing the morphometric values, regions associated with bifurcations were considerably larger than the rest of the artery, thus to avoid skewing results 0.5mm was removed from the end of each arterial segment if that section was part of a bifurcation. The arteries could not be reconstructed in one SS 4-week mouse and one SS 24-week mouse.

### Statistical Analysis

All statistical tests were performed in Graphpad Prism 7. Ultrasound and micro-CT data were plotted across age, and data from each group of animals are reported as the mean ± standard error of the mean. A value of p<0.05 was taken to be significant. The effect of genotype was examined using unpaired t-test, and a paired t-test was used to compare measurements taken from ultrasound and micro-CT methodologies. A one-way ANOVA was performed to determine differences across the various ages.

## Results

### Enlarged common carotid arteries in SS mice at 12 weeks of age measured by ultrasound and micro-CT

From ultrasound imaging, luminal diameters were measured in the common carotid and the extracranial portion of the internal carotid arteries of mice 4, 12, and 24-weeks of age. An average diameter between two points 0.5 mm and 1.5 mm upstream from the carotid bifurcation was calculated for the common carotid artery (Fig 1A). At 12-weeks, CCA luminal diameters in SS mice during systole were 19.3% and 11.5% larger than those of age-matched AS mice on the left and right sides, respectively (n=4, p<0.05) (Fig 1B). The CCA remained significantly enlarged during diastole with a 20.1% and 13.7% diameter increase occurring in the left and right sides of SS mice. In the extracranial ICA, a single point, 0.5 mm distal from the carotid bifurcation was measured, and no significant differences in this segment at any age. Table 2 contains all of the measurements from ultrasound imaging of the extracranial carotid arteries.

**Figure 1:**
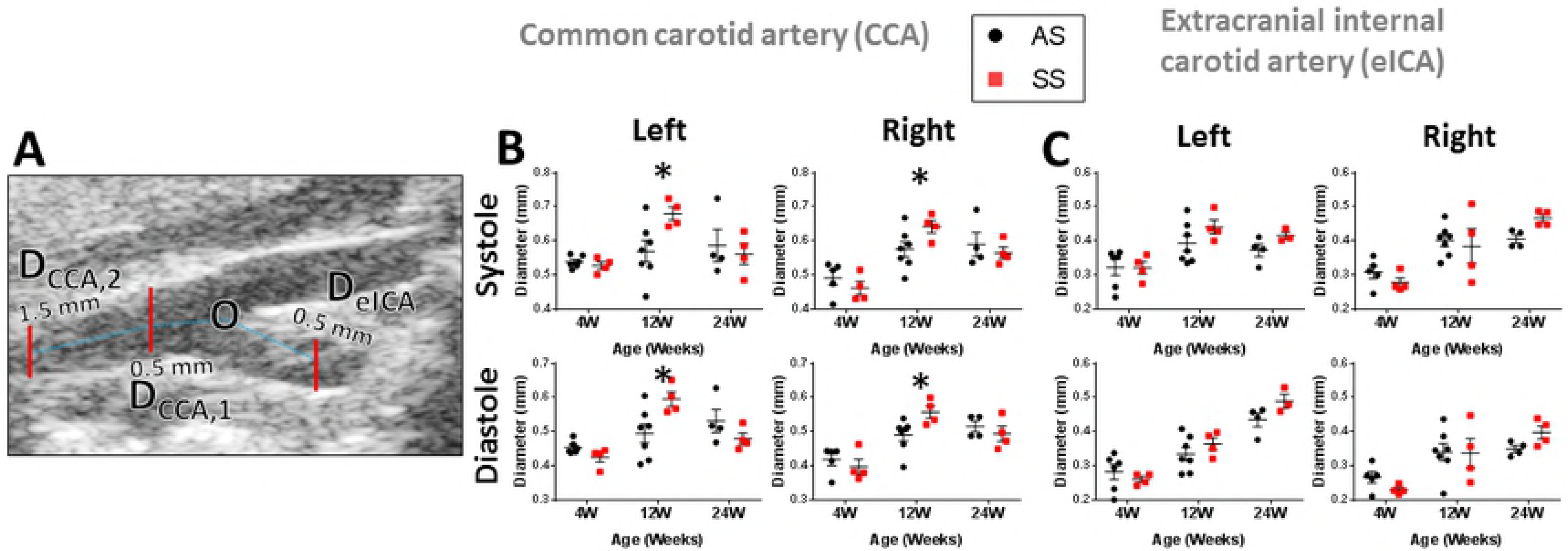
The common carotid artery (CCA) luminal diameter is enlarged in SS mice at 12 weeks of age during the entire cardiac cycle. **(A)** The luminal diameter was measured at in the CCA 0.5 and 1.5 mm upstream from the carotid bifurcation (labeled O), and in eICA 0.5 mm downstream from the carotid bifurcation. **(B)** The common carotid artery was significantly larger in 12-week SS mice during both systole and diastole for the left and right sides. **(C)** There are no significant differences found in the eICA diameters of SS and AS mice. *p < 0.05 SS compared with AS by T-test. Error bars are SEM.

**Table 2.** Morphological measurements from ultrasound of the extracranial internal carotid arteries in mice with sickle trait (AS) and sickle cell anemia (SS).

|  | Systolic |  |  |  |  |  | Diastolic |  |  |  |  |  |
| --- | --- | --- | --- | --- | --- | --- | --- | --- | --- | --- | --- | --- |
| Age (Weeks) | 4 weeks |  | 12 weeks |  | 24 weeks |  | 4 weeks |  | 12 weeks |  | 24 weeks |  |
| Genotype | AS | SS | AS | SS | AS | SS | AS | SS | AS | SS | AS | SS |
| Left CCA Diameter (mm) | 0.539<br>(0.007) | 0.531<br>(0.015) | 0.57<br>(0.031) | 0.68<br>(0.019)<sup>a</sup> | 0.54<br>(0.029) | 0.464<br>(0.007) | 0.454<br>(0.008) | 0.422<br>(0.02) | 0.495<br>(0.027) | 0.596<br>(0.021)<sup>a</sup> | 0.587<br>(0.047) | 0.532<br>(0.034) |
| Right CCA Diameter (mm) | 0.493<br>(0.022) | 0.473<br>(0.024) | 0.576<br>(0.022) | 0.642<br>(0.018) | 0.59<br>(0.035) | 0.576<br>(0.019) | 0.419<br>(0.018) | 0.409<br>(0.027) | 0.491<br>(0.018) | 0.558<br>(0.018)<sup>a</sup> | 0.516<br>(0.016) | 0.493<br>(0.032) |
| Left eICA Diameter (mm) | 0.323<br>(0.023) | 0.309<br>(0.018) | 0.393<br>(0.023) | 0.441<br>(0.021) | 0.435<br>(0.02) | 0.495<br>(0.035) | 0.282<br>(0.022) | 0.257<br>(0.01) | 0.335<br>(0.019) | 0.365<br>(0.018) | 0.373<br>(0.019) | 0.419<br>(0.017) |
| Right eICA Diameter (mm) | 0.311<br>(0.023) | 0.264<br>(0.003) | 0.401<br>(0.017) | 0.385<br>(0.051) | 0.406<br>(0.012) | 0.473<br>(0.013) | 0.268<br>(0.022) | 0.224<br>(0.003) | 0.341<br>(0.025) | 0.338<br>(0.043) | 0.35<br>(0.01) | 0.411<br>(0.019) |

CCA, Common Carotid Artery; eICA, Extracranial Internal Carotid Artery.
<sup>a</sup> $p < 0.05$ compared with AS by t-test. Number in parentheses is standard deviation of the measurement.

Since there was a limitation in obtaining measurements intracranially with ultrasound, we validated that comparable measurements were acquired between ultrasound performed on living mice, and the arterial casts measured by micro-CT, on non-living animals (Fig 2). Mice 12 weeks old were used for comparisons since that was the age when statistically significant differences were observed between SS and AS carotid arteries (Fig 3). Values calculated from the reconstructed arteries were only comparable to systolic ultrasound measurements as a systolic pressure of 100 mmHg was maintained throughout the casting process. The maximum inscribed diameter, which is the largest circle that can fit within the boundaries of a vessel cross-section (Fig 3A), were averaged along the length of the CCA and extracranial ICA. In the CCA this length extended 0.5 to 1.5 mm proximal of the carotid bifurcation, and in the extracranial ICA this extended 0.4 to 0.6 mm downstream of the bifurcation.

**Figure 2:**
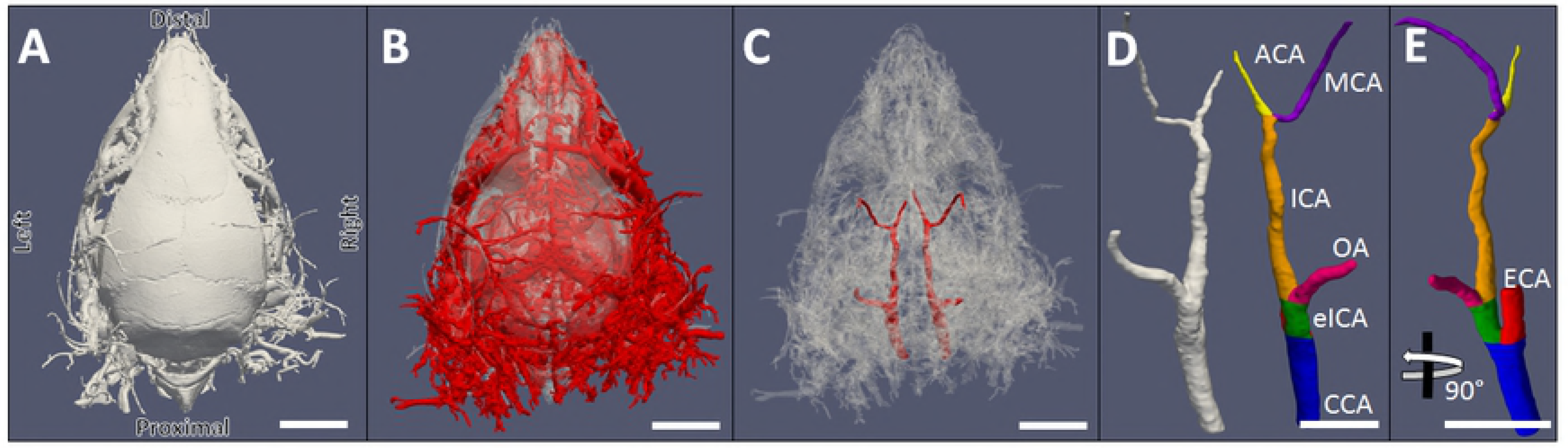
Casts of blood vessels to acquire morphology of carotid and cerebral arteries. Top view of corrosion casting process. **(A)** Microfil, a radio-opaque polymer, was injected into the aorta of mice after exsanguination and fixation of the blood vessels under pressurized conditions. **(B)** The head of mouse was decalcified in a 10% HCL solution for 36 hours to remove bone. **(C)** The reconstructive software Mimics Materialise was used to segment and reconstruct carotid and cerebral arteries. **(D,E)** Final vascular model with individual arterial segments colored. CCA (common carotid artery), eICA (extracranial internal carotid artery), ECA (external carotid artery), OA (ophthalmic artery), intracranial ICA (internal carotid artery), ACA (anterior cerebral artery), MCA (middle cerebral artery). Scale bars = 5 mm.

**Figure 3:**
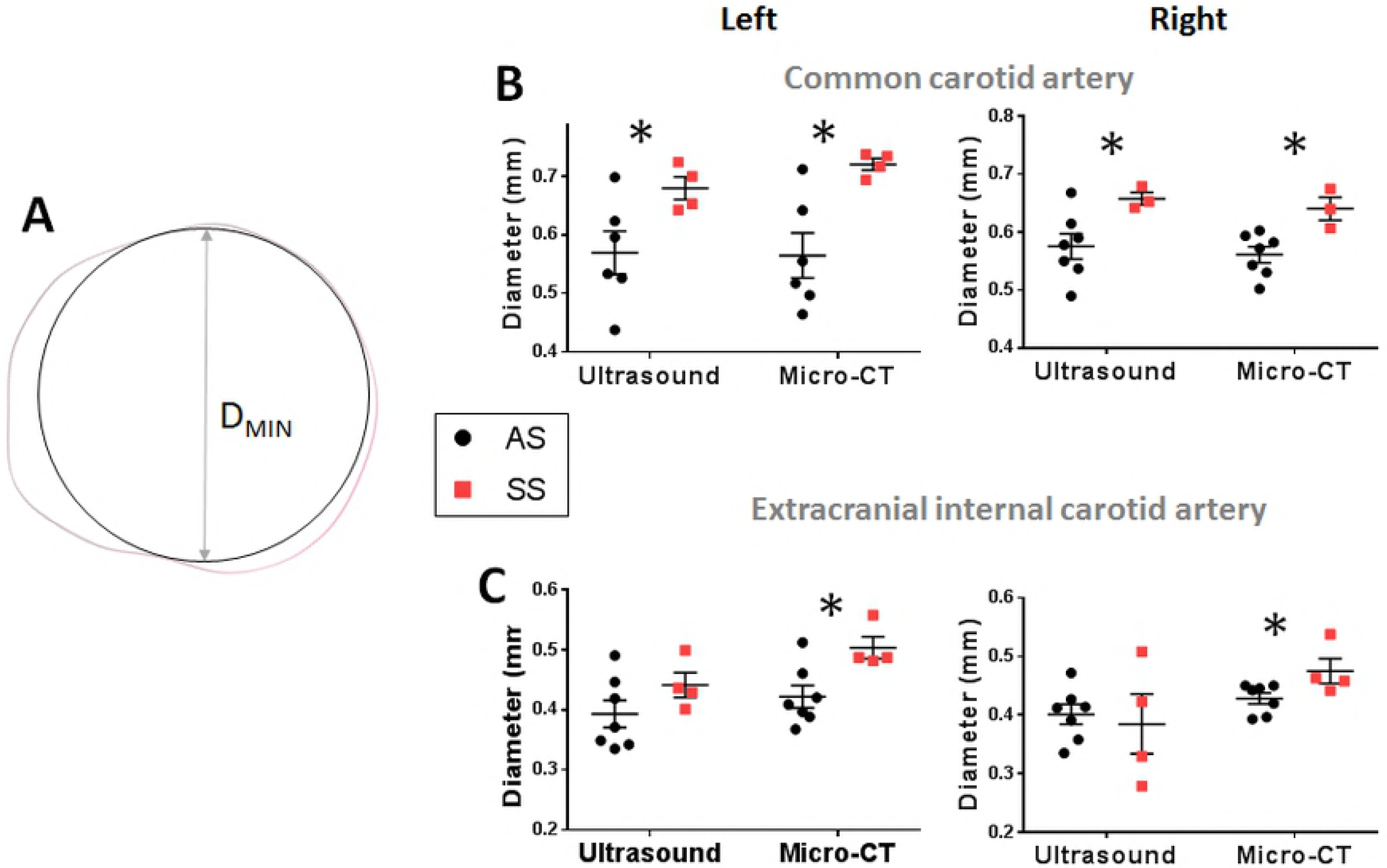
Reconstructed models from artery casts recapitulate ultrasound measurements from living animals. Comparison of diameters measured from ultrasound and micro-CT in the 12-week age group. **(A)** micro-CT measurements reveal CCA diameters were larger in SS mice, paralleling results from the ultrasound study. No significant differences were found between the CCA diameters measured through ultrasound and diameters calculated from reconstructed micro-CT images. **(B)** Diameters calculated from micro-CT were significantly larger in SS mice. No significant differences were observed between both methodologies, however, diameters calculated through micro-CT had smaller SEM. *p<0.05 SS compared with AS by T-test. Paired T-test was performed between ultrasound and μCT methodologies for the same genotype. Error bars are SEM.

There were no significant differences between diameters from ultrasound and micro-CT methodologies (paired T-test, n=4, p>0.05). Diameters calculated through the micro-CT method paralleled the significant differences in the common carotid artery of the 12-week group for SS vs. AS mice (Fig 3B). Statistically significant differences were measured in the extracranial ICA using the micro-CT technique, but not by ultrasound (Fig 3C).

### Diameter vs. perimeter and cross-sectional area as reliable metrics

Diameter measurements were useful to extract data from the ultrasound, but it assumed a perfectly circular geometry, which was not usually the case for these arteries. Using the micro-CT analysis of corrosion casts, cross-sectional shapes of the artery were evaluated for ellipticity, a metric that can be used to determine the roundness of polygons, a value of 0 is perfectly circular, and values closer to 1 indicate a flatter object. The ellipticity in both AS and SS was found to be in the range between 0.4 and 0.6, denoting non-circular geometry (Figure 4A), although the diameters were determined (Figure 4B), and useful for comparing methods of ultrasound vs. micro-CT (Figure 3). Using micro-CT, the cross-sectional perimeter and area of carotid arteries were calculated along the same length of the artery, better capturing the irregular geometry presented by the arteries. The perimeter and cross-sectional area were 34.6% and 65% larger, respectively, in the left extracranial ICA of SS mice 12 weeks of age (p<0.05, n=3), whereas the right extracranial ICA increase was only 8.3% and 18.4%, which was not significant (Fig 4C,D). There were no significant differences in either perimeter or area at 4 or 24 weeks between AS and SS mice. Table 3 contains all of the measurements from micro-CT imaging of corrosion casts for the extracranial carotid arteries.

**Figure 4:**
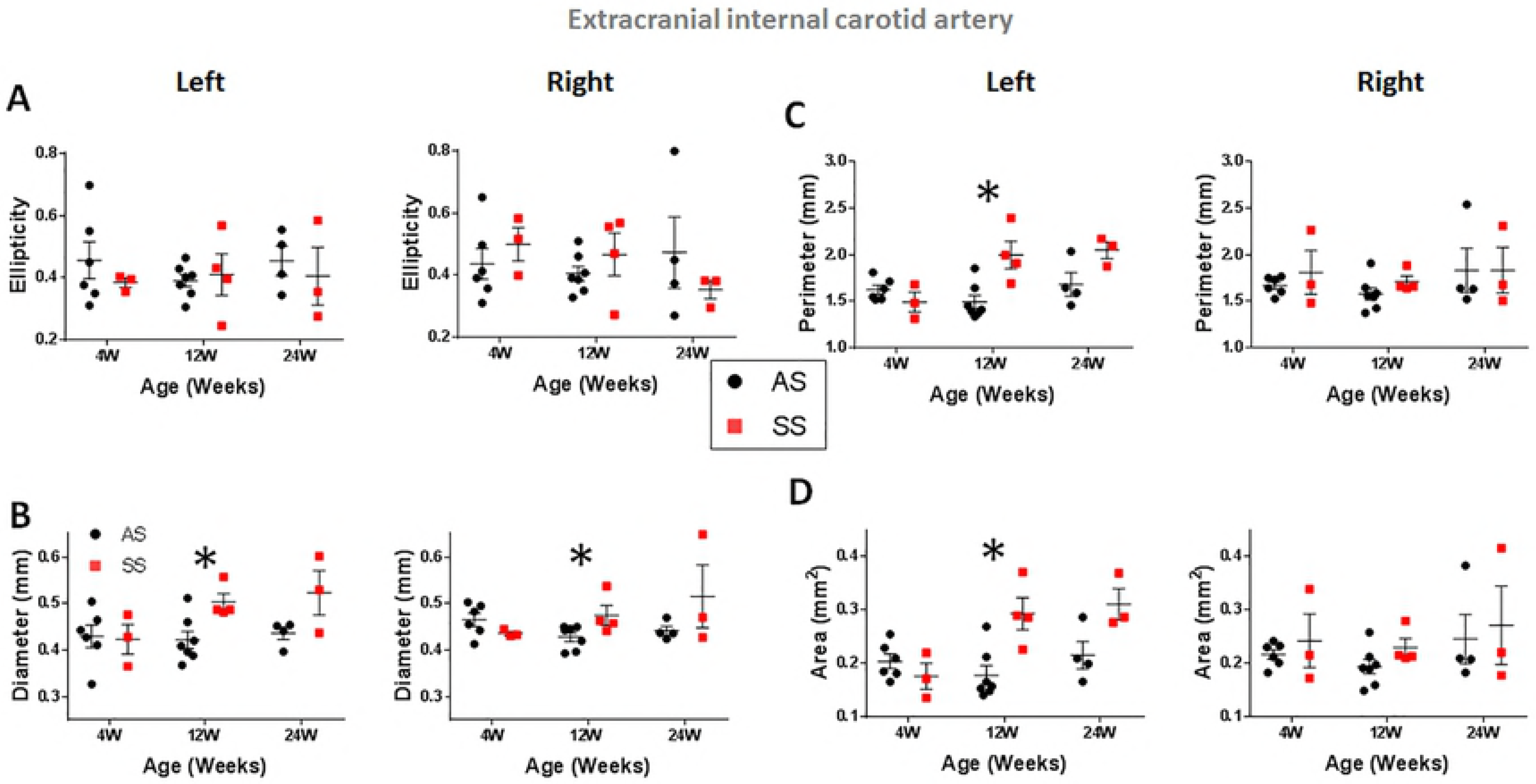
The extracranial internal carotid artery is enlarged on the left side. **(A)** The maximum inscribed circle in a cross-section of the internal carotid artery. **(B)** The maximum inscribed diameter in the extracranial ICA was found to be significantly larger in SS mice for both the left and right sides at 12 weeks. **(C)** The average ellipticity was calculated along the extracranial internal carotid artery from 0.4 - 0.6 mm downstream of the carotid bifurcation. Ellipticity for both AS and SS mice are between 0.4 and 0.6, denoting a more irregular shape (E = 0). Further examination of **(D)** perimeter and **(E)** cross-sectional area reveals only the left side to have significantly larger morphometric values at 12 weeks. *p < 0.05 SS compared with AS by T-test. Error bars are SEM

**Table 3.** Morphological measurements from micro-CT analysis of the extracranial internal carotid arteries in mice with sickle trait (AS) and sickle cell anemia (SS).

|  | Left Extracranial Internal Carotid Artery |  |  |  |  |  | Right Extracranial Internal Carotid Artery |  |  |  |  |  |
| --- | --- | --- | --- | --- | --- | --- | --- | --- | --- | --- | --- | --- |
|  | 4 weeks |  | 12 weeks |  | 24 weeks |  | 4 weeks |  | 12 weeks |  | 24 weeks |  |
|  | AS | SS | AS | SS | AS | SS | AS | SS | AS | SS | AS | SS |
| Diameter (mm) | 0.43<br>(0.024) | 0.423<br>(0.032) | 0.421<br>(0.019) | 0.503<br>(0.018)<sup>a</sup> | 0.436<br>(0.013) | 0.523<br>(0.048) | 0.465<br>(0.014) | 0.436<br>(0.004) | 0.428<br>(0.01) | 0.475<br>(0.021)<sup>a</sup> | 0.442<br>(0.01) | 0.516<br>(0.068) |
| Perimeter (mm) | 1.624<br>(0.048) | 1.491<br>(0.106) | 1.494<br>(0.071) | 1.997<br>(0.147)<sup>a</sup> | 1.681<br>(0.124) | 2.048<br>(0.088) | 1.67<br>(0.039) | 1.809<br>(0.235) | 1.579<br>(0.065) | 1.71<br>(0.058) | 1.833<br>(0.237) | 1.831<br>(0.244) |
| Area (mm<sup>2</sup>) | 0.204<br>(0.014) | 0.176<br>(0.024) | 0.178<br>(0.018) | 0.293<br>(0.03)<sup>a</sup> | 0.216<br>(0.026) | 0.311<br>(0.029) | 0.217<br>(0.009) | 0.243<br>(0.05) | 0.194<br>(0.014) | 0.229<br>(0.017) | 0.246<br>(0.046) | 0.272<br>(0.073) |

eICA, Extracranial Internal Carotid Artery.
<sup>a</sup> $p < 0.05$ compared with AS by T-test.
Number in parentheses is standard deviation of the measurement.

### Internal carotid artery length was similar between SS and AS mice

Internal carotid artery measurements were captured with corrosion casting and micro-CT. When examining the entire ICA (extracranial + intracranial) a significant increase occurred as mice aged from 4 to 12 weeks. Length in the left ICA increased by 11% (p<0.001) and 8% (p<0.01) in AS and SS mice, respectively, with similar increases of 16% (p<0.001) and 11% (p=0.0525) for the right ICA (Fig 5A). There was no significant difference between 12 and 24 weeks. Comparisons of body weight showed similar trends from 4 to 12 weeks. Body weight was 57% (p<0.001) and 53% (p<0.001) greater from 4 to 12 weeks in AS and SS mice, respectively. At 24 weeks, weight increased by 3% (p>0.05) in AS and 17% (p<0.05) in SS. No significant differences were found between genotypes at the same age. These results suggest that radial differences observed in the carotid arteries were not a result of differences in the length or weight of the animal.

**Figure 5:**
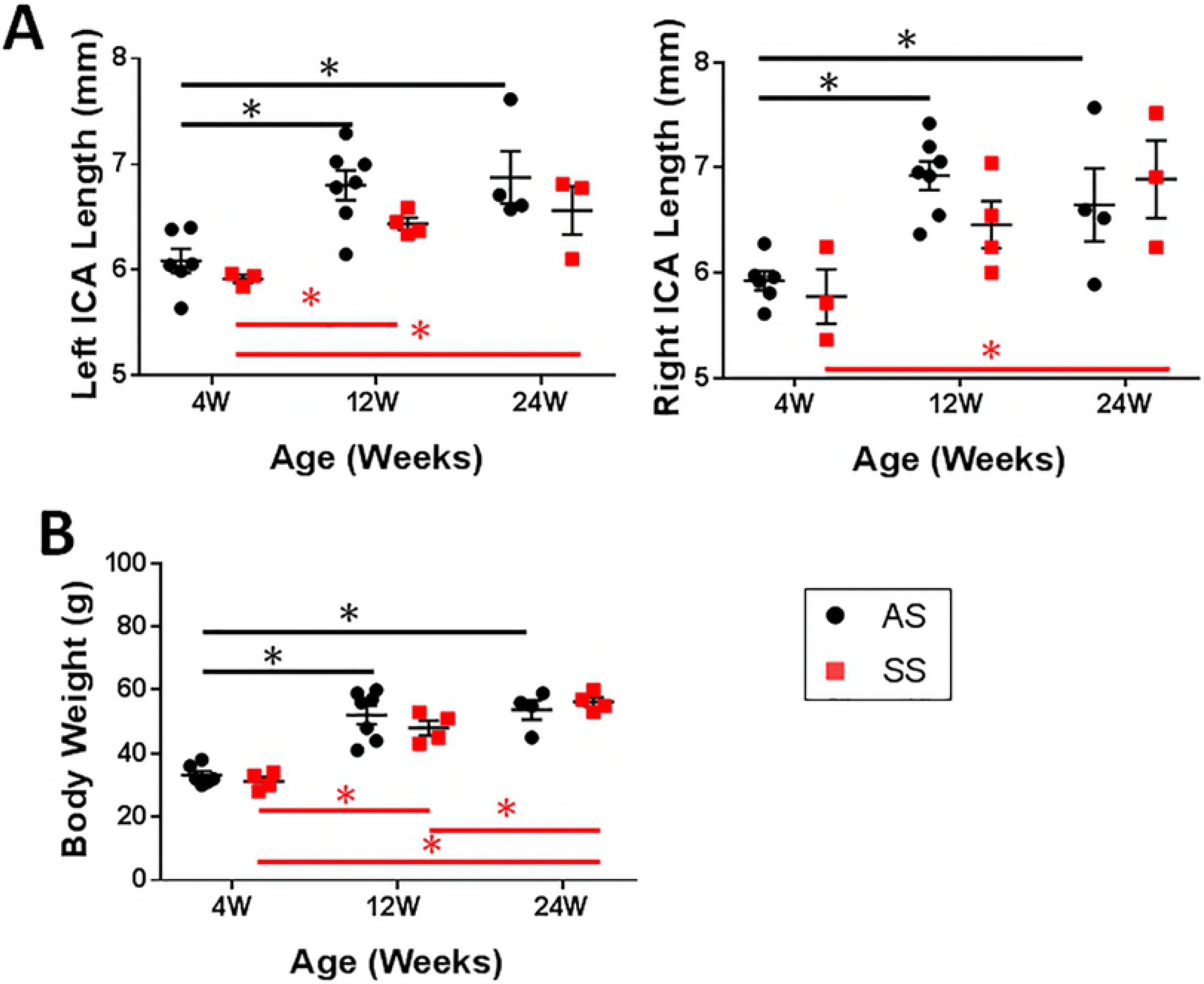
Size and axial growth is similar between homozygous and heterozygous sickle mice. **(A)** Growth axially is quantified based on ICA length (extracranial + intracranial). ICA length increases significantly from 4 to 24 weeks in AS and SS, and no significant differences were found between AS and SS mice in any of the age groups. **(B)** Significant increase in body weight occurs from 4 to 12 weeks in AS and SS mice, but no differences were found in the same age group between genotypes. SS was compared with AS via T-test. The same genotype was compared across ages using a one-way ANOVA, *p < 0.05.

### Narrowing occurs in the intracranial ICA of 12 and 24-week SS mice

Next, we compared intracranial internal carotid arteries (ICA) reconstructed from 4, 12 and 24-weeks old mice. Averaged morphometric values were taken from two sections: (1) the proximal end, which begins 0.5mm from the beginning of the ICA to the midpoint of the vessel; and (2) the distal end, which extends from to midpoint to 0.5 before the end of the vessel (Fig 6A). At 12-weeks, the proximal diameter, perimeter, and cross-sectional area on the left-side of SS mice were enlarged compared to AS controls, increased by 15% (0.048+/-0.02), 16% (1.322 mm +/− 0.025), and 34% (0.139 mm^2^ +/− 0.005), respectively (Fig 6C,D). This difference continued distally, where a 16% (p<0.05) and 32% (p<0.05) increase was marinated in perimeter and area. At 24 weeks, the left proximal section was also enlarged for SS mice, with a 17% increase in perimeter and 40% increase in area as compared to AS. Unlike, their younger counterparts, morphometric values were not significantly different at distal ICA.

**Figure 6:**
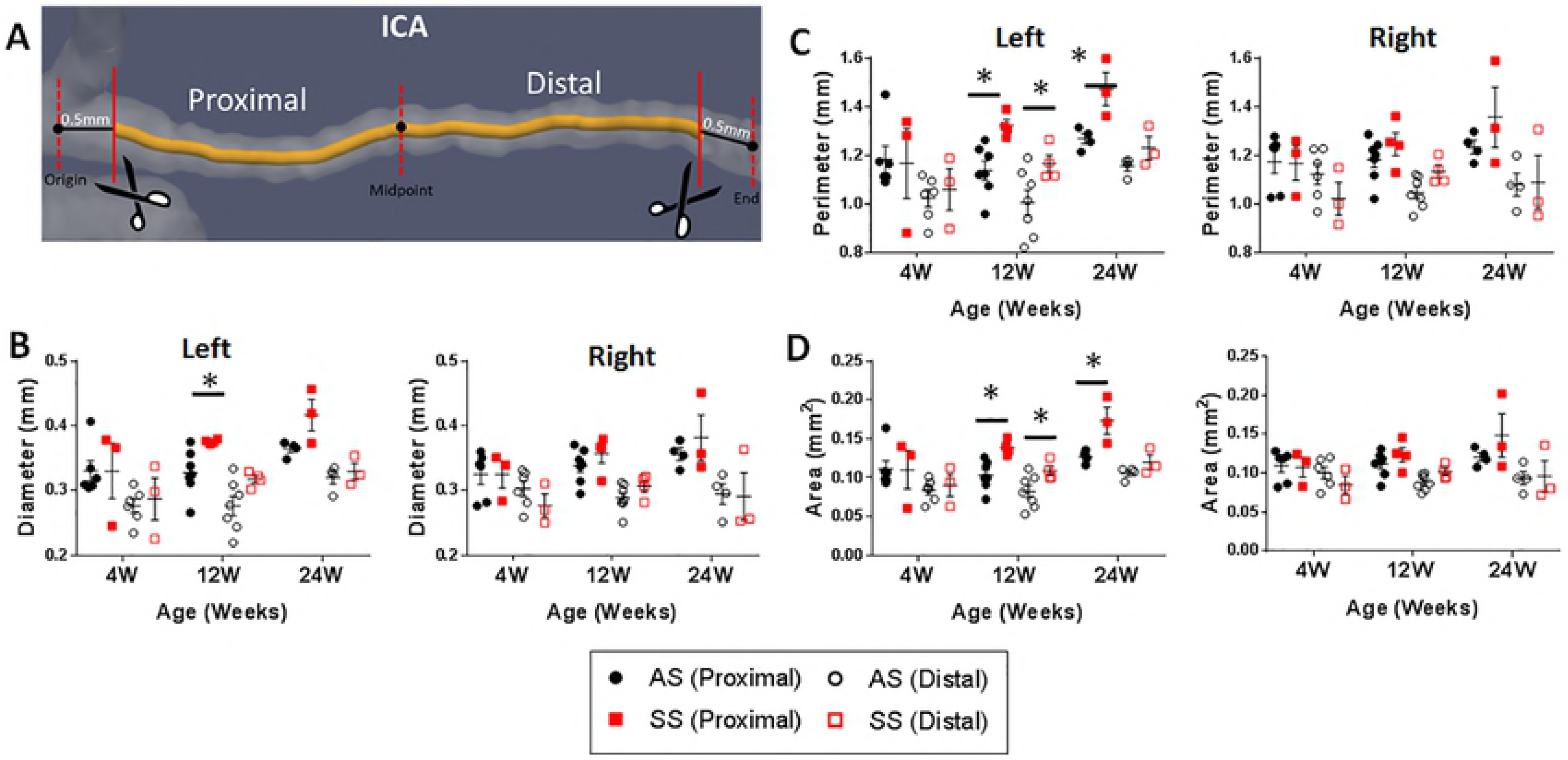
Radial growth of the distal intracranial ICA in SS mice is slowed between 12 and 24 weeks of age. **(A)** The intracranial ICA was divided into two regions: <u>proximal</u>, extending 0.5 from the origin to the midpoint and <u>distal</u> which continues from the midpoint to 0.5 mm before the ICA bifurcation (labeled end). **(B)** Diameter, **(C)** perimeter, and **(D)** cross-sectional area is compared between AS and SS mice at the proximal and distal segments of the intracranial ICA. The left ICA is significantly larger by all morphometric values at 12 weeks at the proximal and distal ends. The proximal ICA is also enlarged in 24-week SS mice but narrows at distal portion losing significance at that end of the ICA. *p<0.05 SS compared with AS by T-test. Error bars are SEM.

These changes from the proximal to distal ICA could be visualized by plotting the mean area for each genotype along the length (Fig 7A). At 4 weeks, SS and AS mice had similar areas along the length. However, in the 12 and 24-week age groups SS mice had significantly larger cross-sections at the proximal ICA, which narrowed, leading to a loss of differences once reaching the opposite end of the artery. To quantify this narrowing in the ICA, area was normalized to the area at the proximal start of the artery for each mouse (Fig 7B). Using this method, variability between specimens can be accounted, and differences previously unseen begin to emerge. The normalized area in 12-week SS mice was as much as 36% lower compared to AS mice. At 24 weeks these differences were even larger; at certain points along the ICA SS mice was 96% and 49% lower in the left and right sides, respectively. Both the left and right intracranial internal carotid arteries narrowed in 12 and 24-week-old SS mice. Table 4 contains the measurements from micro-CT imaging of corrosion casts for the intracranial internal carotid arteries.

**Figure 7:**
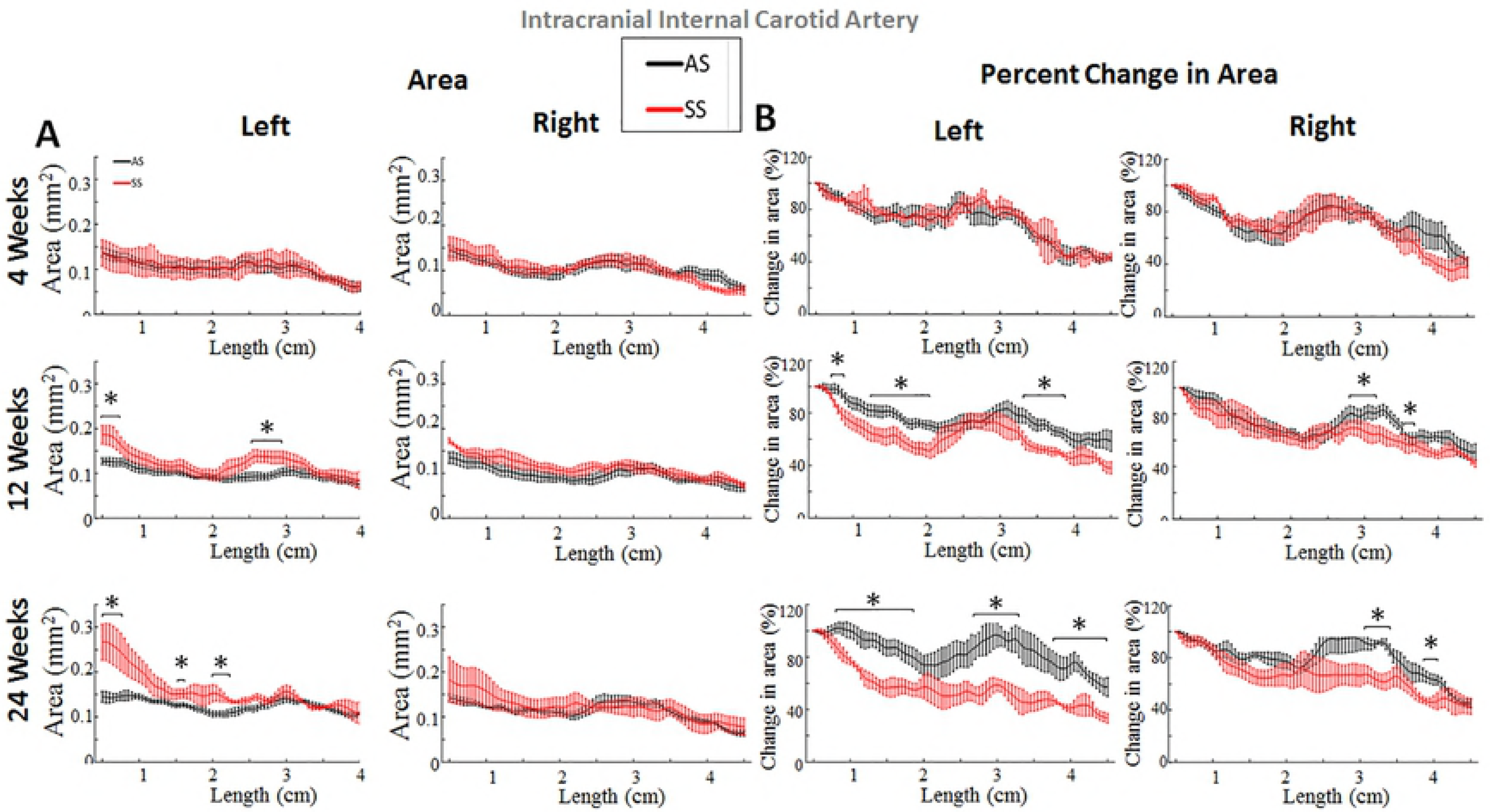
Distal Intracranial Internal Carotid Artery Narrows in 12 and 24-week SS mice. Mean values for bulk area and percent change in area are plotted along the length of the intracranial ICA. **(A)** The left proximal ICA is enlarged in 12-and 24-week SS mice, but differences are lost upon moving towards the distal section of the artery. **(B)** Area was normalized to the starting point of artery to account for variability between subjects. The ICA narrows significantly from the proximal to distal end of the artery in 24-week SS mice as compared to age-matched controls. *p<0.05 SS compared with AS by T-test. Error bars are SEM.

**Table 4.** Morphological measurements from micro-CT analysis of the intracranial internal carotid arteries in mice with sickle trait (AS) and sickle cell anemia (SS).

|  |  | Left ICA |  |  |  |  |  | Right ICA |  |  |  |  |  |
| --- | --- | --- | --- | --- | --- | --- | --- | --- | --- | --- | --- | --- | --- |
|  |  | 4 weeks |  | 12 weeks |  | 24 weeks |  | 4 weeks |  | 12 weeks |  | 24 weeks |  |
|  |  | AS | SS | AS | SS | AS | SS | AS | SS | AS | SS | AS | SS |
| Diameter (mm) | Proximal | 0.332<br>(0.016) | 0.33<br>(0.044) | 0.33<br>(0.013) | 0.377<sup>a</sup><br>(0.003) | 0.365<br>(0.006) | 0.423<br>(0.026)<sup>a</sup> | 0.326<br>(0.014) | 0.326<br>(0.021) | 0.34<br>(0.009) | 0.36<br>(0.015) | 0.355<br>(0.009) | 0.389<br>(0.041) |
|  | Distal | 0.268<br>(0.011) | 0.277<br>(0.03) | 0.272<br>(0.012) | 0.312<sup>a</sup><br>(0.007) | 0.316<br>(0.008) | 0.32<br>(0.016) | 0.289<br>(0.011) | 0.268<br>(0.015) | 0.286<br>(0.008) | 0.305<br>(0.005) | 0.295<br>(0.012) | 0.292<br>(0.035) |
| Perimeter (mm) | Proximal | 1.187<br>(0.054) | 1.169<br>(0.146) | 1.143<br>(0.038) | 1.322<sup>a</sup><br>(0.025) | 1.271<br>(0.022) | 1.49<br>(0.072)<sup>a</sup> | 1.176<br>(0.045) | 1.169<br>(0.071) | 1.186<br>(0.029) | 1.252<br>(0.048) | 1.227<br>(0.026) | 1.368<br>(0.136) |
|  | Distal | 1.012<br>(0.042) | 1.022<br>(0.082) | 0.996<br>(0.044) | 1.152<sup>a</sup><br>(0.039) | 1.141<br>(0.011) | 1.208<br>(0.058) | 1.08<br>(0.039) | 0.268<br>(0.015) | 1.035<br>(0.027) | 1.125<sup>a</sup><br>(0.02) | 1.086<br>(0.041) | 1.1<br>(0.113) |
| Area (mm<sup>2</sup>) | Proximal | 0.112<br>(0.011) | 0.111<br>(0.025) | 0.104<br>(0.007) | 0.139<sup>a</sup><br>(0.005) | 0.128<br>(0.004) | 0.178<br>(0.018)<sup>a</sup> | 0.11<br>(0.008) | 0.108<br>(0.013) | 0.112<br>(0.005) | 0.125<br>(0.009) | 0.119<br>(0.005) | 0.151<br>(0.031) |
|  | Distal | 0.082<br>(0.007) | 0.084<br>(0.013) | 0.08<br>(0.007) | 0.106<sup>a</sup><br>(0.007) | 0.103<br>(0.002) | 0.116<br>(0.011) | 0.093<br>(0.007) | 0.08<br>(0.009) | 0.086<br>(0.004) | 0.1<br>(0.003) | 0.094<br>(0.007) | 0.098<br>(0.02) |
| Length |  | 4.965<br>(0.091) | 4.794<br>(0.068) | 5.587<br>(0.133) | 5.291<br>(0.034) | 5.653<br>(0.196) | 5.29<br>(0.203) | 4.973<br>(0.024) | 4.983<br>(0.15) | 5.724<br>(0.103) | 5.362<br>(0.128) | 5.73<br>(0.283) | 5.744<br>(0.307) |

ICA, Intracranial Internal Carotid Artery.
<sup>a</sup> p < 0.05 compared with AS by T-test.
Number in parentheses is standard deviation of the measurement.

### Portions of the anterior cerebral arteries are significantly narrower in SS mice

Two tributaries branch from the ICA, the middle cerebral and anterior cerebral arteries, and were examined using the same methods from the reconstructed micro-CT models. In the ACA, morphometric values were analyzed along a length of 0.5 to 2.5 mm, whereas the longer MCA calculated values from 0.5 to 4.5 mm of the origin. When examining the bulk area, neither the ACA nor MCA differed significantly in size between AS and SS mice (Fig 8A,B). It should be noted that at 24 weeks, the right ACA in SS mice had significantly smaller diameters (*p<0.05). To investigate these differences further the normalized area was plotted vs. length for the ACA and MCA (Fig 9). Using this approach, the left ACA was found to be enlarged in 4-week SS mice. However, at 12 and 24 weeks the ACA of SS mice were significantly smaller in both the left and right sides, narrowing by up to 60% in some segments. The normalized MCA area between SS and AS mice changed throughout aging. SS mice had enlarged MCAs at 4-weeks, but by 24 weeks the cross-sectional area was narrowed compared to AS controls. Table 5 contains the measurements from micro-CT imaging of corrosion casts for the cerebral arteries.

**Figure 8:**
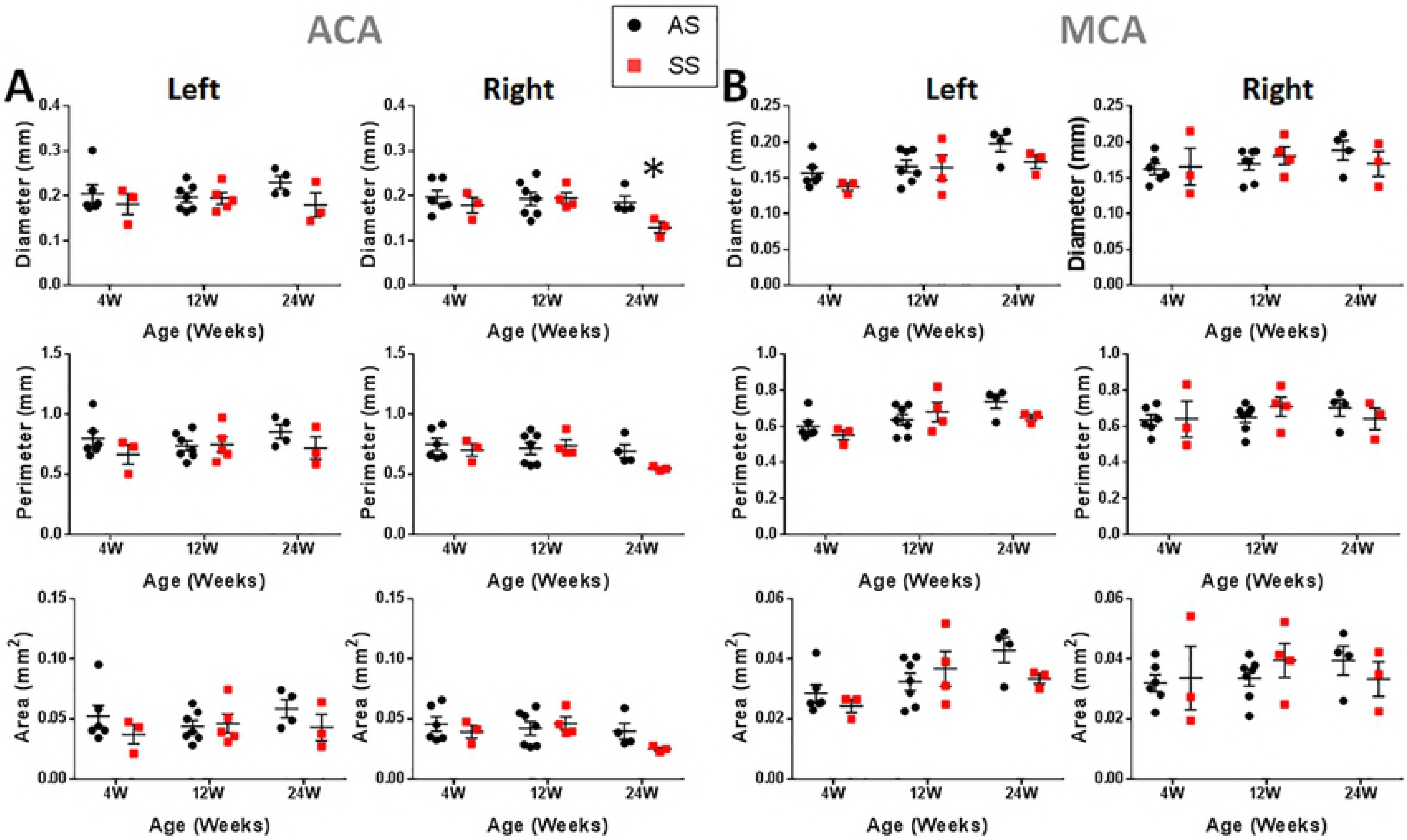
The morphology in the anterior cerebral and middle cerebral arteries are similar between SS and AS mice. **(A)** The ACA does not significantly differ in size between AS and SS mice. At 24 weeks however, the right ACA in SS mice does tend to be smaller with diameters being significantly smaller. **(B)** Morphology in the MCA is not affected by genotype as the diameter, perimeter, and cross-sectional area are not significantly different, regardless of age or side. *p < 0.05 SS compared with AS by T-test. Error bars are SEM.

**Figure 9:**
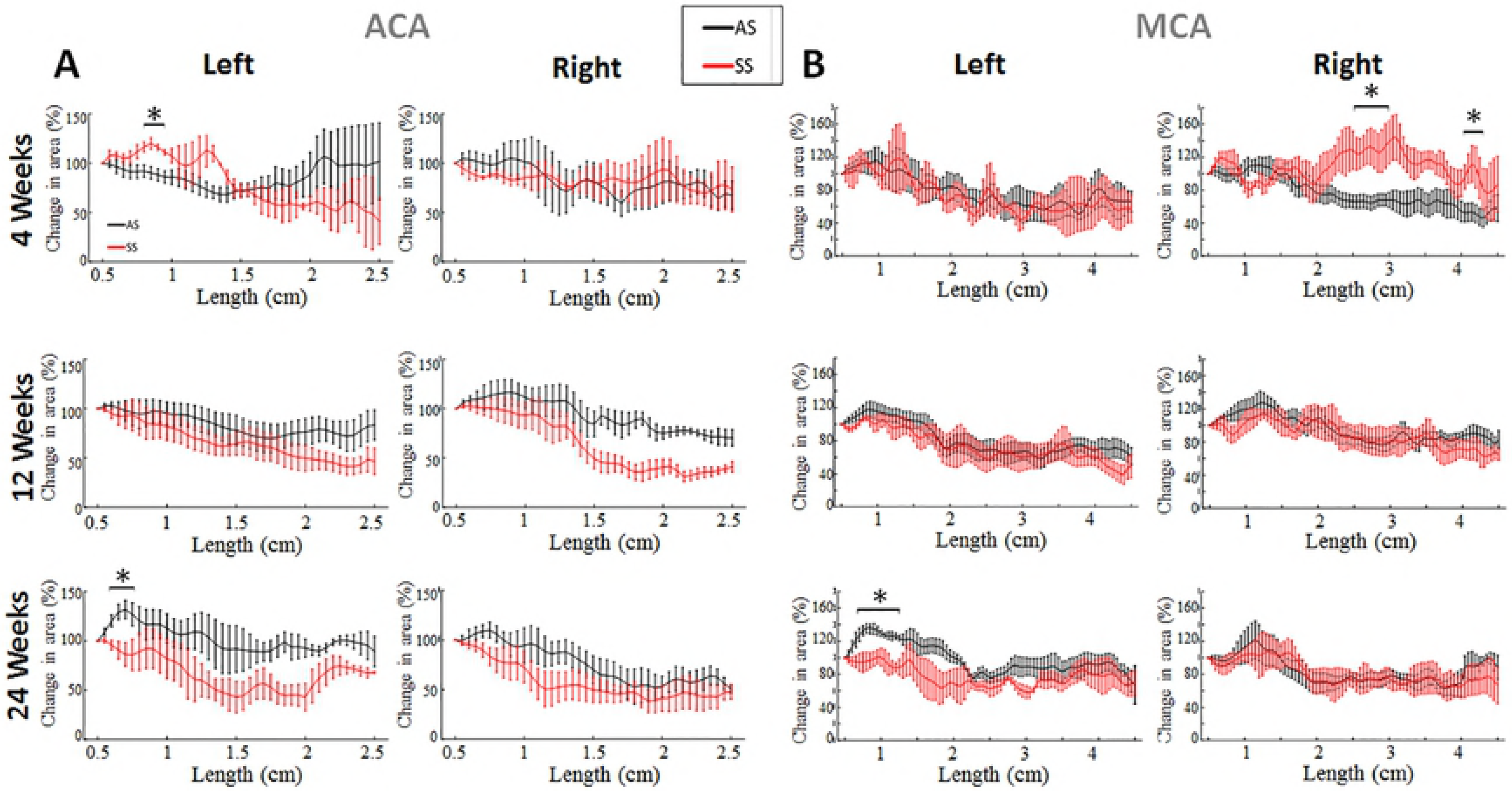
Portions of the anterior cerebral arteries are significantly narrower in SS mice. Mean values of percent change in area are plotted along the length of the ACA and MCA. **(A)** The left ACA is enlarged in 4-week SS mice. At 12 and 24 weeks the ACA of SS mice becomes narrowed at various portions of the ACA, in both the left and right sides. **(B)** The relation in normalized MCA area between SS and AS mice changes throughout age. SS mice have enlarged MCAs at 4-weeks, but at 24 weeks the cross-sectional area is narrowed compared to AS controls. *p < 0.05 SS compared with AS by T-test. Error bars are SEM.

**Table 5.** Morphological measurements from micro-CT analysis of the cerebral arteries in mice with sickle trait (AS) and sickle cell anemia (SS).

| Middle Cerebral Artery (MCA) | Left Middle Cerebral Artery |  |  |  |  |  | Right Middle Cerebral Artery |  |  |  |  |  |
| --- | --- | --- | --- | --- | --- | --- | --- | --- | --- | --- | --- | --- |
|  | 4 weeks |  | 12 weeks |  | 24 weeks |  | 4 weeks |  | 12 weeks |  | 24 weeks |  |
|  | AS | SS | AS | SS | AS | SS | AS | SS | AS | SS | AS | SS |
| Diameter (mm) | 0.147<br>(0.007) | 0.136<br>(0.007) | 0.16<br>(0.011) | 0.155<br>(0.012) | 0.187<br>(0.008) | 0.163<br>(0.006) | 0.158<br>(0.009) | 0.161<br>(0.026) | 0.161<br>(0.007) | 0.171<br>(0.01) | 0.182<br>(0.01) | 0.162<br>(0.018) |
| Perimeter (mm) | 0.61<br>(0.024) | 0.57<br>(0.031) | 0.617<br>(0.037) | 0.647<br>(0.032) | 0.773<br>(0.037) | 0.657<br>(0.027) | 0.682<br>(0.026) | 0.677<br>(0.09) | 0.626<br>(0.024) | 0.673<br>(0.045) | 0.763<br>(0.025) | 0.651<br>(0.074) |
| Area (mm<sup>2</sup>) | 0.031<br>(0.003) | 0.026<br>(0.003) | 0.031<br>(0.003) | 0.033<br>(0.004) | 0.049<br>(0.005) | 0.036<br>(0.004) | 0.039<br>(0.003) | 0.04<br>(0.009) | 0.031<br>(0.002) | 0.036<br>(0.005) | 0.048<br>(0.003) | 0.035<br>(0.008) |
| Anterior Cerebral Artery (ACA) | Left Anterior Cerebral Artery |  |  |  |  |  | Right Anterior Cerebral Artery |  |  |  |  |  |
|  | 4 weeks |  | 12 weeks |  | 24 weeks |  | 4 weeks |  | 12 weeks |  | 24 weeks |  |
|  | AS | SS | AS | SS | AS | SS | AS | SS | AS | SS | AS | SS |
| Diameter (mm) | 0.218<br>(0.028) | 0.177<br>(0.026) | 0.194<br>(0.011) | 0.198<br>(0.015) | 0.218<br>(0.013) | 0.192<br>(0.025) | 0.193<br>(0.012) | 0.185<br>(0.015) | 0.19<br>(0.014) | 0.194<br>(0.015) | 0.186<br>(0.014) | 0.138<br>(0.016) |
| Perimeter (mm) | 0.888<br>(0.104) | 0.756<br>(0.081) | 0.797<br>(0.035) | 0.895<br>(0.07) | 0.88<br>(0.056) | 0.862<br>(0.101) | 0.848<br>(0.052) | 0.754<br>(0.045) | 0.784<br>(0.039) | 0.807<br>(0.061) | 0.787<br>(0.042) | 0.632<br>(0.057) |
| Area (mm<sup>2</sup>) | 0.066<br>(0.017) | 0.054<br>(0.003) | 0.054<br>(0.004) | 0.084<br>(0.024) | 0.061<br>(0.007) | 0.066<br>(0.014) | 0.060<br>(0.008) | 0.045<br>(0.005) | 0.053<br>(0.005) | 0.0556<br>(0.008) | 0.054<br>(0.007) | 0.036<br>(0.008) |

MCA, Middle Cerebral Artery; ACA, Anterior Cerebral Artery. Standard error of the mean is shown in parentheses
<sup>a</sup> p < 0.05 compared with AS by T-test.
Number in parentheses is standard deviation of the measurement.

## Discussion

Here we used a combination of ultrasound and micro-CT imaging to quantify and characterize morphology in the carotid and cerebral arteries at different ages of mice to examine differences between mice with sickle cell anemia (SS) or littermate heterozygous (AS) controls. Morphometric values calculated from micro-CT imaging of extracranial carotid artery casts paralleled measurements from the live mouse during systole. This suggest that morphometric values calculated in the cerebral arteries are within physiological norms. Significant differences found only through micro-CT in the extracranial ICA was a result of the greater resolution at which points can be measured. Through ultrasonography only a few distinct points can be measured in a vessel; however, using the reconstructed arteries multiple points can be analyzed along the length and averaged together. This reduces the overall error in the measurement. Additionally, the diameter measured from ultrasound is of a cross-section, and did not consider the irregular shape of the vessel. Measurements calculated through the reconstructed artery can provide area and perimeter, thereby quantifying the true shape of the vessel.

The greatest differences when examining the bulk morphometric values were observed between AS and SS mice in the 12-week age group. The left common carotid artery diameter was larger in SS mice as measured through both ultrasound and micro-CT measurements. This result agrees with findings obtained by Cahill et. al, who compared the homozygous sickle Townes mice to C57BL/6J wild-type counterparts in 10 to 13-week-old mice [17] and also found SS mice to have significantly larger common carotid arteries measured via ultrasound.

Extending downstream, the entire intracranial ICA was significantly larger on the left side in SS mice 12 weeks of age. At 24 weeks however, only the proximal portion of the ICA was found to have significantly larger cross-sectional perimeters and areas. These results parallel findings in humans, as direct measurements via magnetic resonance angiography have shown luminal area to be larger in the ICA, ACA, and MCA of patients with SCD as compared to age matched controls [18]. Additionally, although there was a significant growth from 4 to 12 weeks; axially, ICA length was similar between SS and AS mice, and overall body weight showed no difference, matching previous studies [19]. Therefore, the radial differences observed in SS mice, is not due to differences in animal size but is a result of the animal genotype.

In the distal intracranial ICA, however, a distinct narrowing was observed which led to a loss of the statistically significant difference between SS and AS mice at 24-weeks of age. This narrowing was quantified by normalizing the area to the most proximal point of the artery and plotting the percent change in area against length. Through this method we can see that the ICA was significantly narrower at certain portions of the distal ICA, in right and left sides of 12 and 24-week week old mice. Using this method again, significant narrowing occurred in the ACA of SS mice (Fig 9), despite seeing no significant differences in the bulk morphometric properties, except for the ACA diameter being smaller in 24-week SS mice. As previously stated, high velocities occur in cerebral arteries with sickle cell anemia, and are associated with stroke risk [4-6]. The narrowing luminal area in SS mice suggests a potential for hemodynamic changes as well.

Differences between AS and SS were mostly observed in the 12-week age group. Age is known to be a strong factor of stroke risk in sickle cell anemia, with the highest risk of ischemic stroke occurring between the ages of two and five in humans [1]. Ages of mice do not correlate linearly to human age, as maturation in the first month of life occurs 150 times faster and later months have a third of the previous maturation rate [20], however, some approximations can be made to compare with human lifespan and marking clinical pathologies of humans with sickle cell anemia with findings in the mouse model. At four weeks, mice are in early adolescence (approximating humans of 10-12 years old), while at 12 weeks mice approximate humans in their early 20s. At 24 weeks, the mice approximate humans in their early 30s. Individuals with SCA in their 20s have an elevated risk of hemorrhagic stroke [2]; the bulk morphometric differences observed in 12-week mice may be related to this type of stoke. Though no aneurysms were found in the mice themselves, uncontrolled expansion of the artery can lead to hemorrhagic stroke. At 24 weeks of age, mice approximate the age of humans in their 30s, and this is the age at which humans with SCA have an elevated risk of ischemic stroke. The narrowing observed in the cerebral arteries of mice with SCA could potentially cause a stroke as blockages in these narrowed sections of the artery may prevent an adequate supply of blood and oxygen.

The left common carotid and internal carotid artery lumens were larger in SS mice compared to AS mice. The cause for these differences is not fully known. It has been previously reported that the hippocampus, thalamus, and visual cortex on the left side of the brain in C57BL/6J mice, one of the background strains for the Townes mice, are enlarged [21]. These regions are supplied by several arteries that branch from the ICA and may have higher oxygen demands than their counterparts on the right. This could potentially be magnified in a mouse with sickle cell anemia, leading to the larger artery size that was observed.

In conclusion, this study characterizes morphological characteristics in the carotid and cerebral arteries of SS and AS mice from ages 4 – 24 weeks. Casting of the arteries corresponds to *in vivo* measurements and provides information in regions of the vasculature that cannot be easily imaged with other modalities in mice. Morphological differences between genotypes were found to occur in 12-and 24-week mice, coinciding with the different types of strokes that are most prevalent in humans with SCA. This suggests that SCA affects the vascular morphology, continually changing the cerebrovascular system through the entire lifespan of the subject, whether mouse or human. Through quantification of the morphometric values it may be possible to predict the type and likelihood of stroke in an individual with SCA. Future work is needed to determine the cause of these morphological changes, and ultimate consequences they produce.

## References

1. Hillery CA, Panepinto JA. Pathophysiology of stroke in sickle cell disease. Microcirculation. 2004;11(2):195–208.

2. Ohene-Frempong K, Weiner SJ, Sleeper LA, Miller ST, Embury S, Moohr JW, et al. Cerebrovascular accidents in sickle cell disease: rates and risk factors. Blood. 1998;91(1):288–94.

3. Merkel KH, Ginsberg PL, Parker JC, Post MJ. Cerebrovascular disease in sickle cell anemia: a clinical, pathological and radiological correlation. Stroke. 1978;9(1):45–52.

4. Riela AR, Roach ES. Etiology of stroke in children. J Child Neurol. 1993;8(3):201–20.

5. Switzer JA, Hess DC, Nichols FT, Adams RJ. Pathophysiology and treatment of stroke in sickle-cell disease: present and future. Lancet Neurol. 2006;5(6):501–12.

6. Boros L, Thomas C, Weiner WJ. Large cerebral vessel disease in sickle cell anaemia. J Neurol Neurosurg Psychiatry. 1976;39(12):1236–9.

7. Okuyama S, Okuyama J, Okuyama J, Tamatsu Y, Shimada K, Hoshi H, et al. The arterial circle of Willis of the mouse helps to decipher secrets of cerebral vascular accidents in the human. Med Hypotheses. 2004;63(6):997–1009.

8. Ryan TM, Ciavatta DJ, Townes TM. Knockout-transgenic mouse model of sickle cell disease. Science. 1997;278(5339):873–6.

9. Hsu LL, Champion HC, Campbell-Lee SA, Bivalacqua TJ, Manci EA, Diwan BA, et al. Hemolysis in sickle cell mice causes pulmonary hypertension due to global impairment in nitric oxide bioavailability. Blood. 2007;109(7):3088–98.

10. Aslan M, Ryan TM, Adler B, Townes TM, Parks DA, Thompson JA, et al. Oxygen radical inhibition of nitric oxide-dependent vascular function in sickle cell disease. Proc Natl Acad Sci U S A. 2001;98(26):15215–20.

11. Duvall CL, Taylor WR, Weiss D, Guldberg RE. Quantitative microcomputed tomography analysis of collateral vessel development after ischemic injury. Am J Physiol-Heart C. 2004;287(1):H302–H10.

12. Abruzzo T, Tumialan L, Chaalala C, Kim S, Guldberg RE, Lin A, et al. Microscopic computed tomography imaging of the cerebral circulation in mice: Feasibility and pitfalls. Synapse. 2008;62(8):557–65.

13. Huo Y, Guo X, Kassab GS. The flow field along the entire length of mouse aorta and primary branches. Ann Biomed Eng. 2008;36(5):685–99.

14. Yanagisawa H, Davis EC, Starcher BC, Ouchi T, Yanagisawa M, Richardson JA, et al. Fibulin-5 is an elastin-binding protein essential for elastic fibre development in vivo. Nature. 2002;415(6868):168–71.

15. Mattson DL. Comparison of arterial blood pressure in different strains of mice. Am J Hypertens. 2001;14(5 Pt 1):405–8.

16. Antiga LS, S. VMTK: Vascular Modeling Toolkit. http://wwwvmtkorg/. 2009.

17. Cahill LS, Gazdzinski LM, Tsui AKY, Zhou YQ, Portnoy S, Liu E, et al. Functional and anatomical evidence of cerebral tissue hypoxia in young sickle cell anemia mice. J Cerebr Blood F Met. 2017;37(3):994–1005.

18. Vaclavu L, Baldew ZAV, Gevers S, Mutsaerts H, Fijnvandraat K, Cnossen MH, et al. Intracranial 4D flow magnetic resonance imaging reveals altered haemodynamics in sickle cell disease. Br J Haematol. 2018;180(3):432–42.

19. Xiao L, Andemariam B, Taxel P, Adams DJ, Zempsky WT, Dorcelus V, et al. Loss of Bone in Sickle Cell Trait and Sickle Cell Disease Female Mice Is Associated With Reduced IGF-1 in Bone and Serum. Endocrinology. 2016;157(8):3036–46.

20. Fox JG. The mouse in biomedical research. 2nd ed. Amsterdam; Boston: Elsevier, AP; 2007. v. <2–4> p.

21. Spring S, Lerch JP, Wetzel MK, Evans AC, Henkelman RM. Cerebral asymmetries in 12-week-old C57Bl/6J mice measured by magnetic resonance imaging. Neuroimage. 2010;50(2):409–15.

